# Environmental impacts of Broiler chicken production in the North Eastern Himalayan region of India: Evaluation using the Life Cycle Assessment approach

**DOI:** 10.64898/2026.03.16.712132

**Authors:** Mahak Singh, Trinayana Kaushik, P. L. Bhutia, Rekha Yadav, Vinay Singh, Rahul Katiyar, H. Kalita

## Abstract

The study presents the first comprehensive environmental assessment of broiler chicken production in India, from a cradle-to-farm-gate perspective, using the Life Cycle Assessment (LCA) approach. The objective of the study was to identify environmental *hotspots* in the broiler production system. All inflows and outflows of the broiler production system were mapped to construct the life-cycle inventory. This study uses a system boundary that extends from the cradle to the farm gate. Inventory data for the broiler chicken farms were collected from six different farms, which is typical of the Indian broiler production system, and background data were sourced from the Ecoinvent 3.0 database. For the LCA study, SimaPro (v 9.3.0.3) was used with the ReCiPe 2016 Midpoint impact assessment methodology. For this study, the functional unit of the “1-kg live weight chicken produced was taken into consideration”. The environmental impact categories assessed were mainly Global Warming Potential in hundred years (GWP), stratospheric ozone depletion, freshwater eutrophication, terrestrial ecotoxicity, land use, etc. Results showed that broiler chicken feed was primarily responsible for environmental impacts, followed by transportation and electricity. Broiler chicken production had a total GWP of 3.77 kg CO2-eq per kilogram of live weight. Specifically, the energy component of feed, viz., maize production, was the main source of environmental impact. The process of transporting feed and chicks to the broiler farm also had a significant environmental impact. The broiler production system was found to have moderate environmental impacts compared to other published LCA studies of chicken production. Based on the findings of the present study, we proposed actionable items to further improve the environmental efficiency of broiler chicken production in India.

## 1. Introduction

In the years ahead, demand for livestock products is expected to rise dramatically due to rising wealth, urbanization, and a projected global population of 9.6 billion by 2050. The livestock sector is crucial to sustainable food security and health. Globally, poultry meat is the most consumed meat (OECD-FAO, 2020). To keep up with rising demand, global poultry meat production rose dramatically from 9 to 133 million tonnes between 1961 and 2020. In 2020, chicken meat share was 40 percent of total world meat production, with much of the growth observed in Asia, where production has increased almost fourfold (FAOSTAT, 2022).

India is one of the world’s largest producers of eggs and broiler meat. The poultry sector is the fastest-growing agri-sector in India and contributes one percent to the national GDP. Poultry meat accounts for 50 percent of total meat production in India and ranks 5th globally (Government of India, 2019). Chicken meat is widely accepted in India without any religious or socio-cultural restrictions. The substantial growth in per capita income and the urban population, particularly the middle class, has significantly driven demand growth (Singh et al., 2022). The broiler industry has been growing at an average of 8-10 percent over the last 20 years and currently accounts for around two-thirds of total poultry output. The industry has a high level of vertical integration. Integrators supply the chicks to the broiler farm. The broiler farms rear the chicks to slaughter weight. Adult birds are either purchased by a wholesaler or sold directly by broiler farms. Approximately 90-95 percent of total sales volume is live birds, while cold storage accounts for around 5 percent.

On the other hand, the increased chicken meat production contributes significantly to the environmental impacts, including air, land, soil, water, and biodiversity (Costantini et al., 2021; Ogino et al., 2021). There is a need to produce more with limited natural resources and with fewer environmental impacts. Along with milk, poultry products are considered to have the least environmental impact among animal-source foods, especially in terms of carbon footprint and resource depletion. The land use and energy requirement per unit of chicken meat are low (de Vries and de Boer, 2010; Roma et al., 2015). However, when compared with a broader basket of edible fresh foods, these products fall into the medium-high impact category (Clune et al., 2017; Duarte da Silva Lima et al., 2019). Also, there is increased consumer awareness of food with a lower environmental impact.

Life Cycle Assessment (LCA) involves the “compilation and evaluation of the inputs, outputs and potential environmental impacts of a supply chain throughout its life cycle” (ISO 14040, 2006a). In essence, it considers all environmental consequences of resource use, land use, and emissions associated with the product’s processes. Using LCA enables us to assess the environmental impact of products throughout their life cycles, from beginning to end. ISO 14040:2006a and 14044:2006b regulate and standardize LCA, which is founded on four main components (goal and scope definition, inventory, impact assessment, and interpretation of results). LCA is the most widely used methodology for quantifying the environmental impacts of food products (Roy et al., 2009). LCA has long been used to evaluate the environmental impact of food items. Furthermore, it is also a tool for making environmental management decisions (Andersson et al., 1994; Costantini et al., 2021; Djekic and Tomasevic, 2016; Vázquez-Rowe et al., 2012). Life Cycle Assessment of chicken meat production system has been reported from USA (Leinonen et al., 2012; Pelletier et al., 2014; Putman et al., 2017), Canada (Pelletier, 2018), Europe (Cesari et al., 2017; González-García et al., 2014; Tallentire et al., 2019), Australia (Bengtsson and Seddon, 2013; Wiedemann et al., 2017) and Brazil (Duarte da Silva Lima et al., 2019; Prudêncio da Silva et al., 2014). Many LCA studies have been reported on the poultry sector across different regions of North America and the USA, but few have been reported from India. Despite the size and fast growth rate of the broiler industry in India, no reports are available on the environmental efficiency of its production system using LCA. Hence, the current investigation was undertaken with the following objectives: (i) to evaluate the environmental impacts of the broiler production system in India from cradle to farm gate, using the LCA, and (ii) to identify the hot-spots that contribute to environmental impacts in broiler production systems. The findings of this study will be helpful to academics and policymakers in India in addressing the critical environmental impacts of broiler production systems.

## 2. Materials and methods

The study was conducted in Nagaland and Assam, which are the North Eastern states of India (Figure 1). Primary data were collected from six broiler farms representative of the broiler production system in India. The description of the broiler farm and data modeling methodology is described in the following sections.

**Figure 1:**
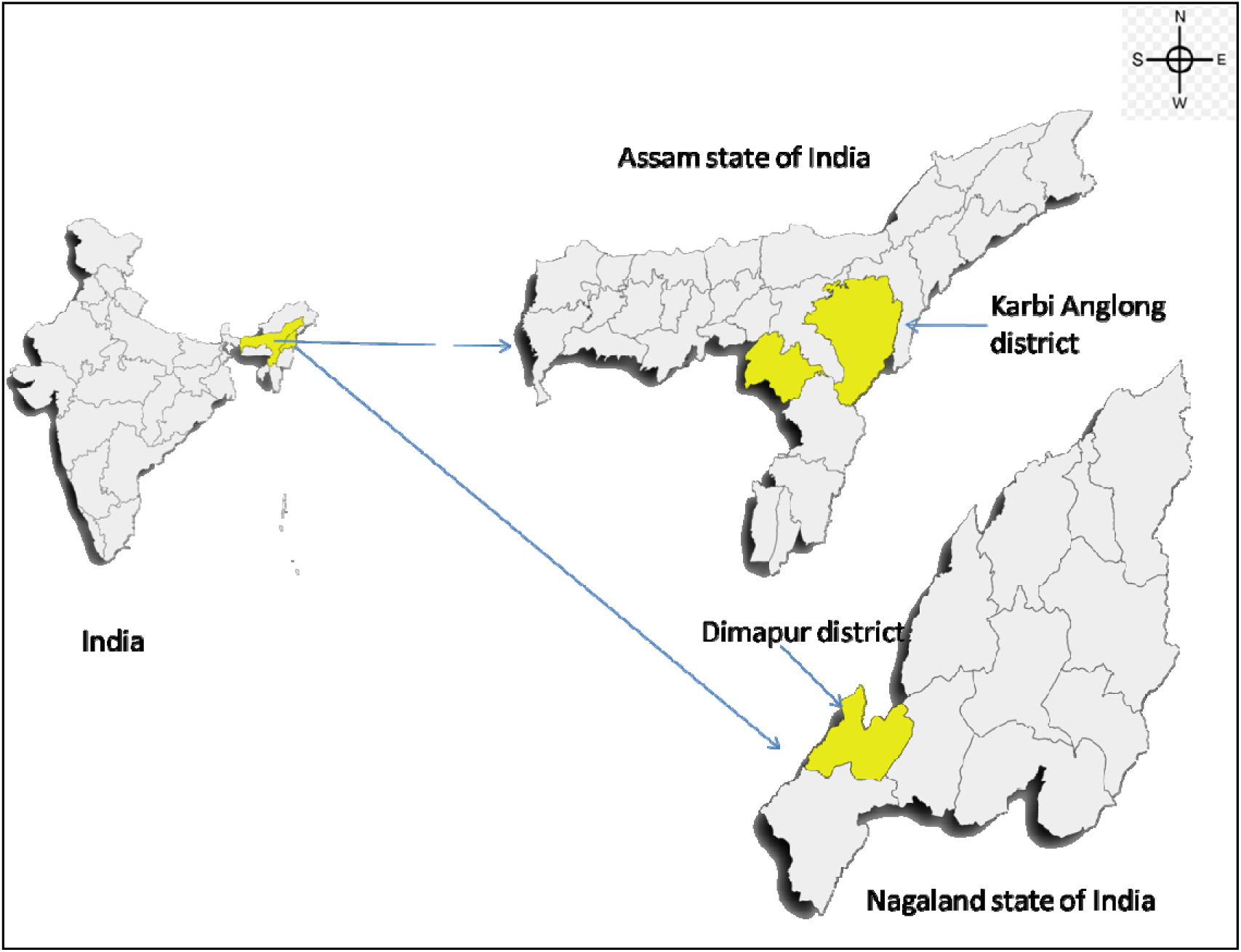
Location of the study area in India (Karbi Anglong district of Assam state and Dimapur district of Nagaland state)

### 2.1. The broiler production system of India

The study site is located in a sub-tropical region (Figure 2) with a monsoon-type climate (Agromet observatory, ICAR Nagaland Centre). In summer, the climate is hot and humid, and in winter it is pleasant and cold. The annual precipitation was 1563.1 mm (mostly from June to September).

**Figure 2:**
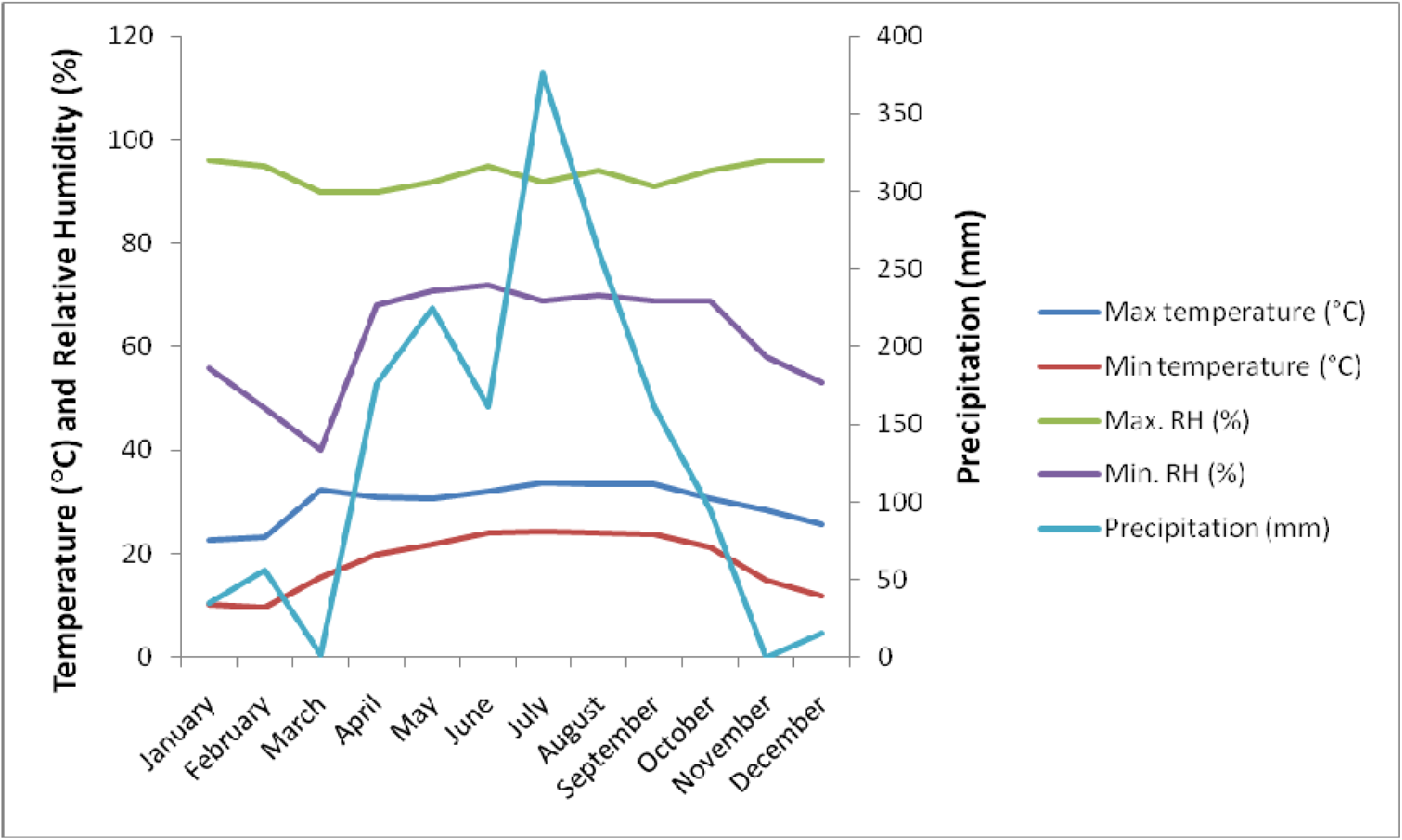
Climate conditions of the study region

The present study analyzed the intensive broiler production system (Figure 3), and for this, primary data (Table 1 and Table 2) were collected from six broiler farms located in Dimapur district of Nagaland (latitude: 25°54′N, Longitude: 93°44′E) and Karbi Anglong district of Assam (latitude: 26.18°N, Longitude:93.58°E). In brief, the broiler production system in India is vertically integrated, with integrators or poultry firms supplying day-old chicks, feed, and medicine to small- to medium-sized contract broiler production farms. After 25 to 42 days, the integrators purchase back the live adult chicken for processing or sale to wholesalers. However, the farmers also raise the broiler chicken independently and sell them to wholesalers or retailers.

**Figure. 3:**
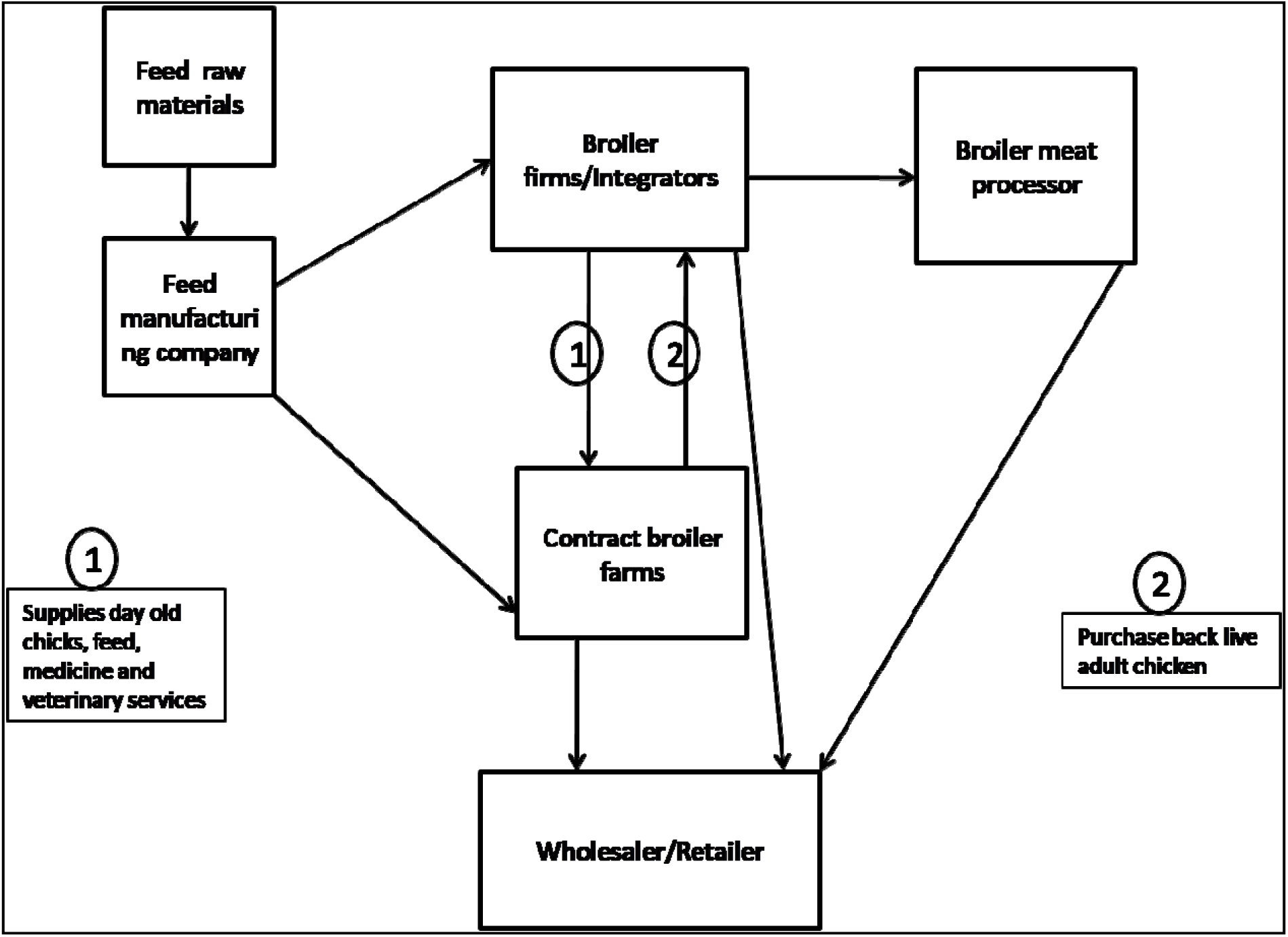
A schematic representation of the broiler production system in India

**Table 1:**
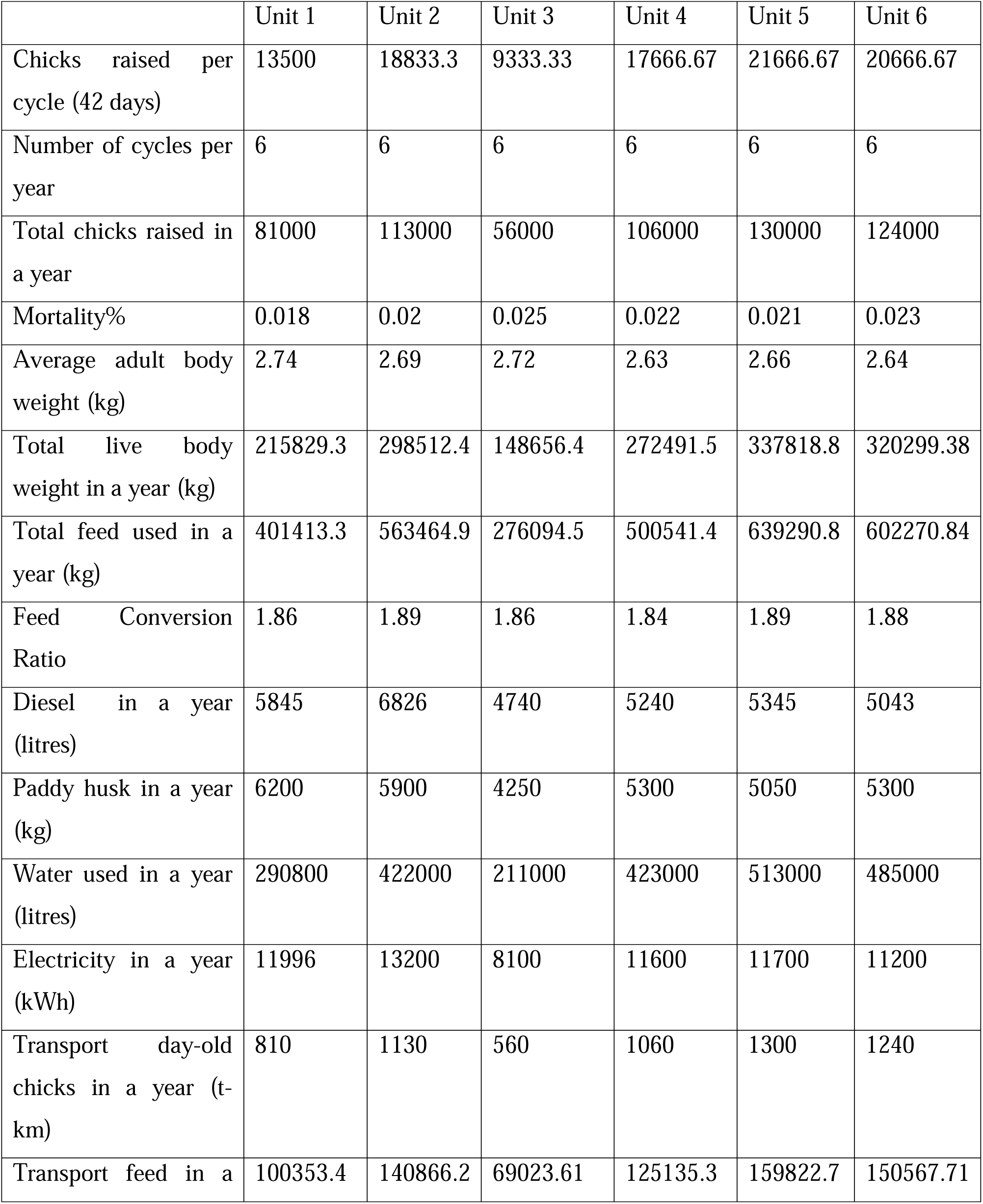
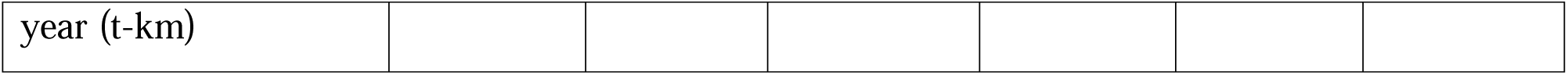
Inventory data of the six different broiler production facilities.

**Table 2:** Average input used for production of one kg live weight broiler chicken#.

| Input | Amount used per kg live weight produced |
| --- | --- |
| Feed (kg) | 1.87 |
| Diesel (litres) | 0.02146 |
| Paddy husk (kg) | 0.02079 |
| Water (litres) | 1.523 |
| Electricity kWh) | 0.44047 |
| Transport day-old chicks (t-km) | 0.00396 |
| Transport (t-km) | 0.4845 |

The study was based on primary data collected from six commercial deep-litter broiler farms, each monitored over six consecutive production cycles within one year (36 cycles in total). All farms were using similar housing design, broiler genetics, veterinary protocols, and feed formulations. These farms are typical of intensive commercial broiler production systems that dominate organized poultry production in India and NEH India and are thus considered representative of modern, integrator-based broiler production in India. We do not aim to represent backyard or smallholder systems, but rather the industrial segment that supplies the majority of formal market broiler meat in the country.

### 2.2. Life Cycle Assessment methodology

LCA is a holistic tool for analyzing the environmental efficiency of a product and includes the environmental impacts of all resources used throughout the product’s life cycle, from cradle to grave (ISO 14040, 2006a; 14044, 2006b; LEAP, 2016). FAO (2016) introduced a harmonized international approach to analyze the environmental performance of poultry supply chains across a wide range of production systems. This study followed the attributional LCA (Alca) approach, i.e., the environmental impacts of the physical inputs and the output flow of the system were assessed. The LCA procedure consists of four phases: (1) goal and scope definition, (2) life cycle inventory analysis (LCI), (3) life cycle impact assessment (LCIA), and (4) life cycle interpretation (Figure 4) (ISO 14040:2006a and 14044:2006b). As per the specification, the broiler production system in the present study is mapped for one year.

**Figure 4:**
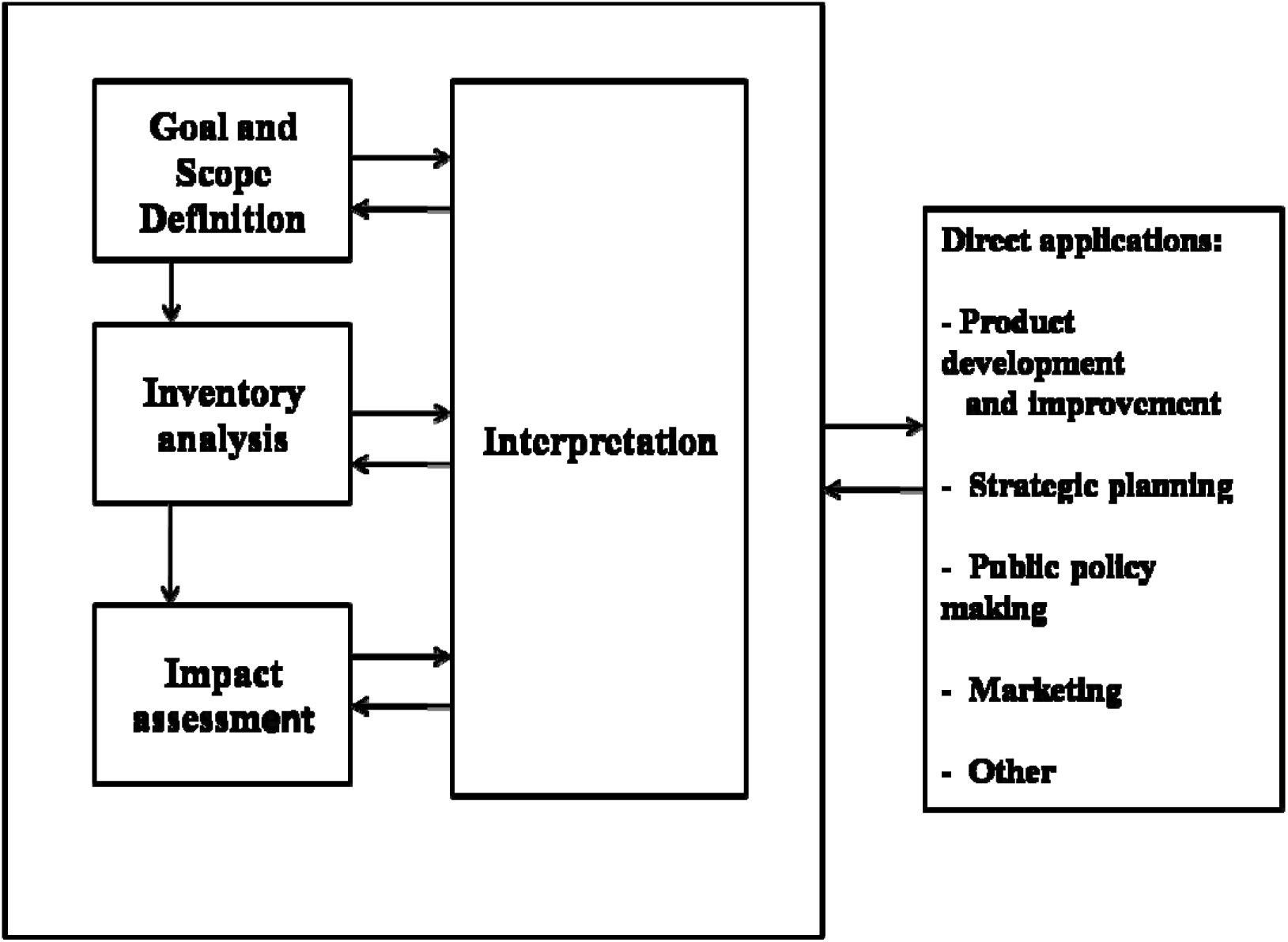
Methodological framework of LCA: phases of an LCA (source: ISO14040, 1997E)

#### 2.2.1. Goal and scope definition

The scope of this study was to analyze the environmental impacts of broiler production systems from day-old chicks to slaughter age (42 days), with a particular focus on quantifying GHG emissions and other impact categories. The time limit of the primary inventory was from day-old chicks to broilers for slaughter (cradle-to-gate studies, ISO 2006a, 2006b).

##### 2.2.1.1. System boundary and functional unit

The system boundaries included in the LCA assessment are depicted in Figure 5. The system boundary started from the transportation of day-old chicks to the broiler production farm. The transportation of chicks to the broiler farm is included in the system boundaries. The chicks are raised in a poultry bed made of paddy husk, which is also included in the system boundary. During the 42-day fattening period, the chickens are fed with starter and finisher compounded feed, mainly comprising maize, palm oil, soya meal, fish meal, limestone, and other ingredients (Table 3). The life cycle inventory for each ingredient was modelled separately and aggregated to obtain the environmental profile of 1 tonne of complete feed, which was then multiplied by the feed intake per kg live weight in each farm cycle. The production and transportation of chicken feed are included in the system boundaries. Throughout the growth phase of chickens, water (for cleaning and drinking), elctricity and diesel were used and are included in the system boundaries. The use of vaccines and antibiotics in raising broiler chickens was excluded from the system boundaries due to insufficient information. Chemicals and detergents were used for cleaning activities on the farm; however, they were not considered for assessment due to their insignificant contribution to environmental impacts, as reported in previous studies (Castanheira et al., 2010; González-García et al., 2014). In India, broiler production systems extend from the cradle to the farm gate (live birds) with minimal slaughterhouse processing. As reported earlier (Costantini et al., 2021), the production phase has the most significant impact on animal products. According to ISO 14044:2006 b, the functional unit shall be consistent with the goal and scope of the study, as well as with the system boundaries. The primary functional unit (FU) was 1 kg live weight (LW) of broiler at the farm gate. In addition, a secondary nutritional functional unit was used: 100 g of edible protein from broiler meat. This allows comparison of environmental impacts per unit mass and per unit of nutritional value.

**Figure 5:**
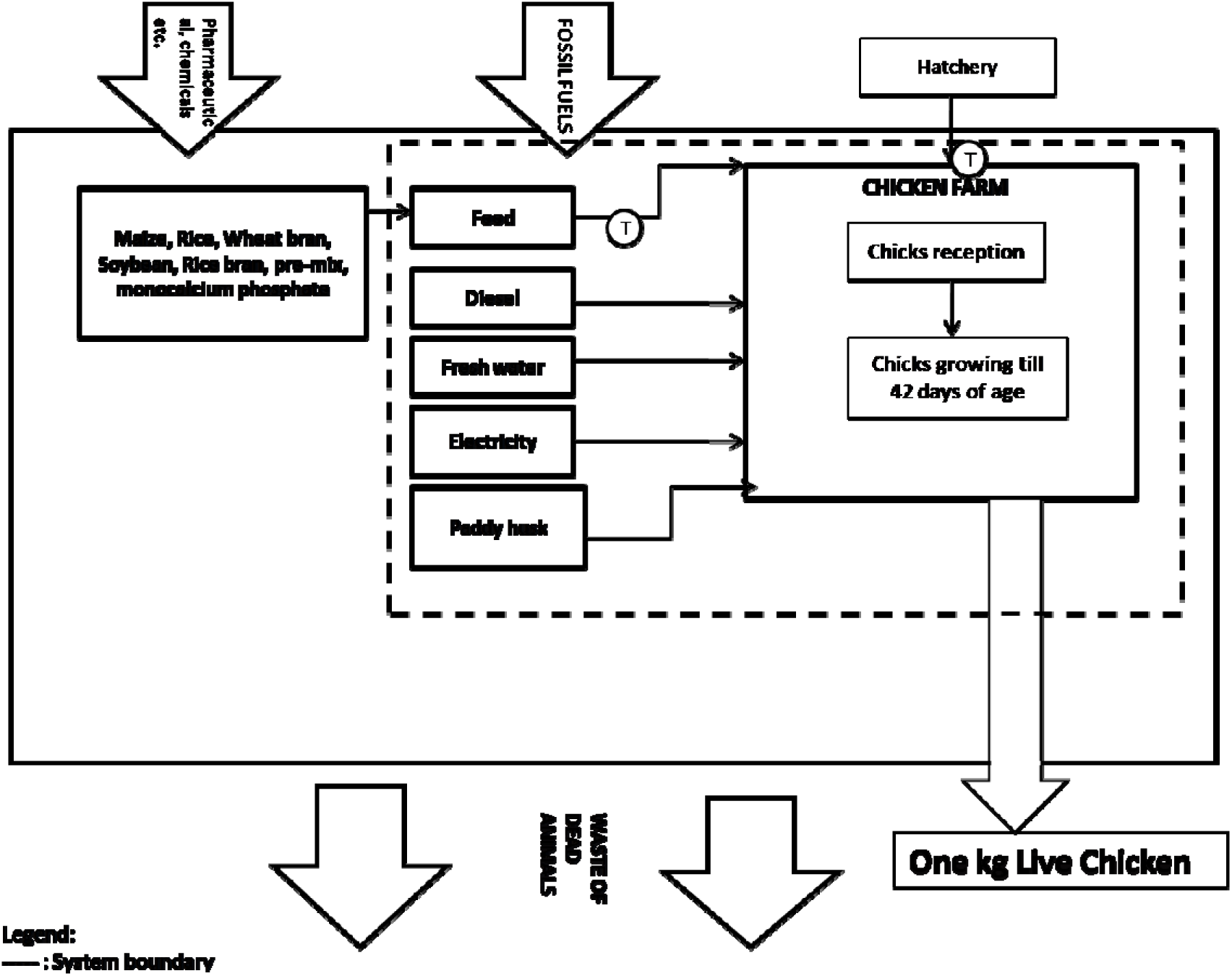
System boundaries and flow diagram of the broiler production system under Life Cycle Assessment

**Table 3:** Aggregated feed composition data for broiler chicken production farm.

| **Ingredients %** | **For 1000 kg of feed** | **For 1.87 kg of feed** |
| --- | --- | --- |
| Maize | 603.40 | 1.13 |
| Palm Oil | 20 | 0.037 |
| Soybean Meal | 292.0 | 0.55 |
| Fish Meal | 50.0 | 0.093 |
| Limestone | 10.0 | 0.018 |
| Di Calcium Phosphate | 12.0 | 0.022 |
| Methionine | 1.0 | .008 |
| Salt | 3.60 | .006 |
| Trace Mineral Mix* | 1.50 | 0.0028 |
| Vitamin premix** | 1.50 | 0.0028 |
| Toxin binder | 5.0 | 0.0093 |
| **Total (kg)** | **1000** | **1.87** |

Birds were reared on deep litter year-round. Litter and manure were not removed between cycles, but accumulated on the floor and were removed once per year and applied to nearby cropland as organic fertilizer. Nitrogen excretion was estimated from feed nitrogen intake minus nitrogen retained in live weight gain, using standard retention coefficients for broilers. Volatile solids (VS) excretion was estimated from feed digestibility assumptions. Methane and nitrous oxide emissions from housing and storage were calculated following IPCC 2019 Refinement Tier 1/2 guidance for poultry, using system-specific emission factors for deep litter. Direct N O emissions were calculated as Nex × EF□, and indirect N□O as (NH□ + NOx volatilization) × EF□. The resulting CH□ and N□O emissions were expressed as CO□-equivalents using the IPCC AR5 100-year GWP values.

Because all collected manure was applied to cropland, we used a substitution approach, assuming that the nutrient content (N, P□O□, K□O) of poultry manure replaced an equivalent amount of synthetic mineral fertilizer. The avoided production and application of synthetic fertilizers were credited as negative emissions in the system, providing a more complete representation of the environmental consequences of the manure management strategy.

###### Cut-off criteria

The following cut-off criteria were selected as per the goal and scope of the study: the production and maintenance of boiler farms (buildings) and equipment were not included in the assessment. Similarly, cleaning products, disinfectants, pharmaceuticals, or chick production (breeding and hatching) were excluded from the system boundary. Due to the significant uncertainty associated with land-use change (LUC), this aspect was not included in the study.

###### Allocation procedures

Allocation is the partitioning of the input or output flows of a process or a product system between the product system under study and one or more other product systems (ISO 14044:2006b). Allocation is used for multi-functional processes and can significantly affect results. As per the literature (Duarte da Silva Lima et al., 2019; González-García et al., 2014), product allocation was not applied in this study. In the broiler chicken production farm, the main product is live chickens. Poultry manure is removed from the poultry farm at the end of a year of production and used as fertilizer. This transaction represents a negligible fraction of the farm’s total output volume. The manure was considered as residual flow without allocation of impacts.

### 2.3. Life cycle inventory analysis (LCI)

In life cycle assessment studies, it is important to have real, reliable data to obtain representative, relevant results. For this study, life cycle inventory data were collected through personal visits to six broiler chicken production farms in the Indian states of Nagaland and Assam in 2022. The broiler chicken farms were comprehensively assessed and inventoried to obtain representative data. Primary data, such as chicks raised per cycle, number of cycles per year, mortality (%), average body weight (kg), total feed used, feed conversion ratio, diesel used in a year, paddy husk used in a year, etc., were collected from the broiler production facilities. For each farm–cycle combination, resource use was normalized per kg live weight and then averaged across cycles and farms. *2.4. Life cycle impact assessment (LCIA)*

The LCIA step involves calculating the environmental impacts of products or systems based on inventory data. As per the international guidelines of LCA methodology (ISO 14040, 2006a), only classification and characterization were undertaken in this study. As per the scope and goal of the study, normalization and weighting were not done. For this study, the impact assessment methodology was based on Life Cycle Impact Assessment (LCAI) methods (ISO 14040, 2006a). The Sima Pro software (v 9.3.0.3) was used to analyze the data. The impact assessment methods used were ReCiPe 2016 Midpoint (H) V1.03 / World (2010) H. The midpoint characterization was selected because it has a stronger relationship with environmental flows and relatively low uncertainty (Hauschild and Huijbregts, 2015).

The broiler production system utilizes resources and emits substances into the environment, affecting air, land, and water quality. These emissions and resource usage are categorized according to different environmental impact categories (ISO 14040, 2006a). The impact categories used in the present study are listed in Table 4. These impact categories were assessed for this study because (1) these are the most widely used and accepted categories in environmental impact studies across the different livestock production systems and (2) environmental impact from livestock production systems is related to these impact categories (Prudêncio da Silva, 2014; LEAP, 2016; Wiedemann et al., 2017).

**Table 4:** The impact categories and their units assessed in this study.

| Impact category | Measurement Units |
| --- | --- |
| Global warming potential | kg CO<sub>2</sub>-eq |
| Stratospheric ozone depletion | kg CFC11-eq |
| Ionizing radiation | kBqCo-60-eq |
| Ozone formation, human health | Kg NO<sub>x</sub>-eq |
| Fine particulate matter formation | kg PM2.5-eq |
| Ozone formation, terrestrial ecosystem | kg NO<sub>x</sub>-eq |
| Terrestrial acidification | kg SO<sub>2</sub>-eq |
| Freshwater eutrophication | kg P-eq-eq. |
| Marine eutrophication | kg N-eq |
| Terrestrial ecotoxicity | kg 1,4-DCB |
| Freshwater ecotoxicity | kg 1,4-DCB |
| Marine ecotoxicity | kg 1,4-DCB |
| Human carcinogenic toxicity | kg 1,4-DCB |
| Land use | m<sup>2</sup>a crop-eq |
| Mineral resource scarcity | kg Cu-eq |
| Fossil resource scarcity | kg oil-eq |
| Water consumption | m<sup>3</sup> |

### 2.5. Life cycle interpretation

The explanation of the inventory phases, results, and impact categories was conducted in accordance with the study’s goal and scope (FAO, 2016).

## 3. Results

The GWP of the broiler production system was found to be 3.77 kg CO_2_-eq per kg of live weight broiler chicken produced (Table 5). The major contributors to the GWP (Figure 6) were feed (2.1 kg CO_2_-eq per kg of live weight), followed by transportation (0.93 kg CO_2_-eq per kg of live weight) and electricity consumption (0.60 kg CO_2_-eq per kg of live weight). In percentage, feed contributed 55.7% of GWP (Figure 6), followed by transportation (24.74%) and electricity (15.91%). The impact of paddy husk and diesel usage on the GWP of broiler chicken production was found to be insignificant. Terrestrial acidification was found to be contributed primarily by feed, followed by transportation and electricity (Figure 7). However, transportation (53%) contributed the maximum to terrestrial ecotoxicity, followed by feed (40.83%). In the case of land use, 96% was contributed by feed. Feed and transportation accounted for 35.77% and 28.44% of fossil resource scarcity, while diesel and electricity accounted for 20.18% and 14.67%, respectively.

**Figure 6:**
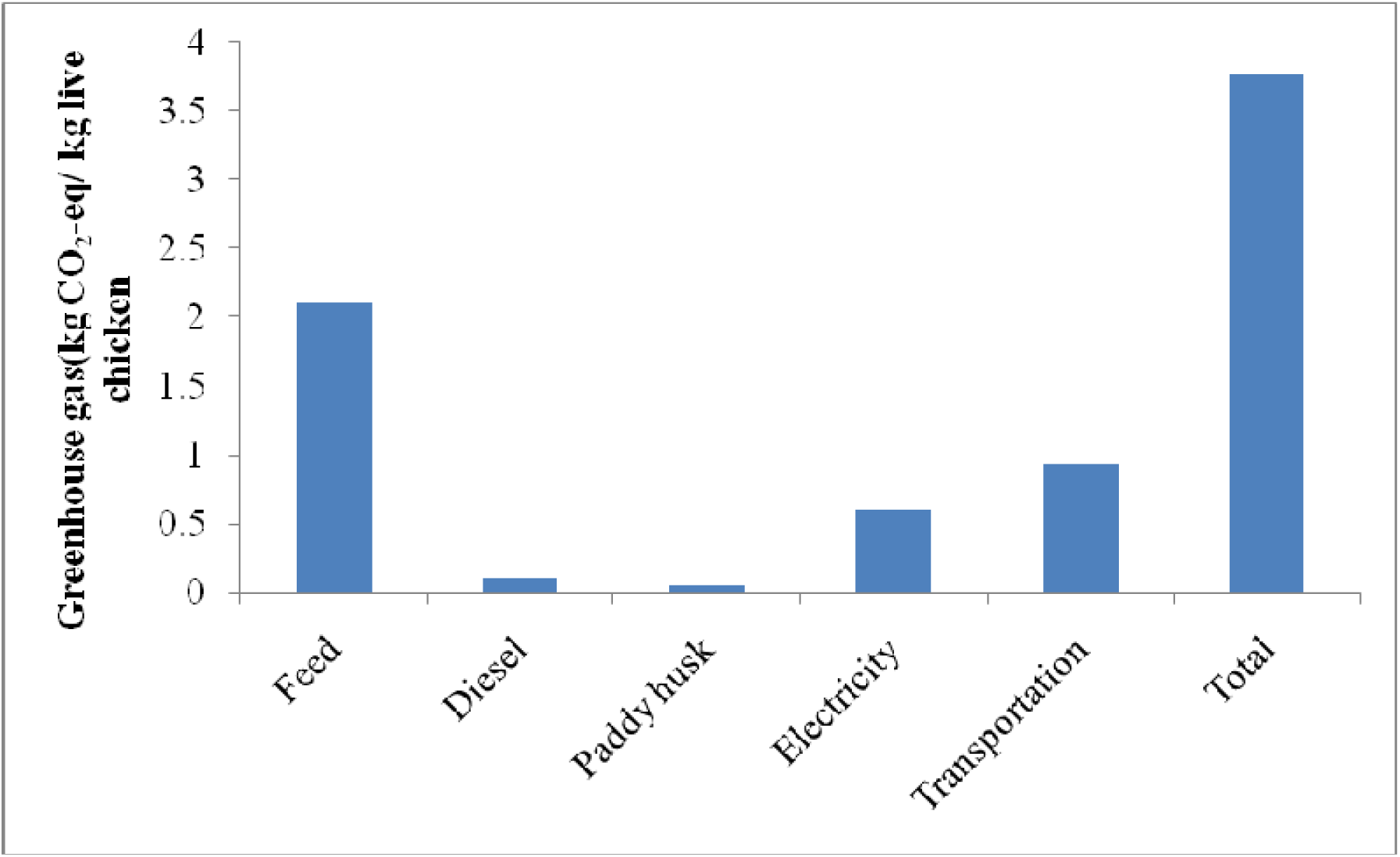
Contribution to greenhouse gas (kg CO_2_-eq/ kg live chicken) emission by different components in broiler chicken meat production in India

**Figure 7:**
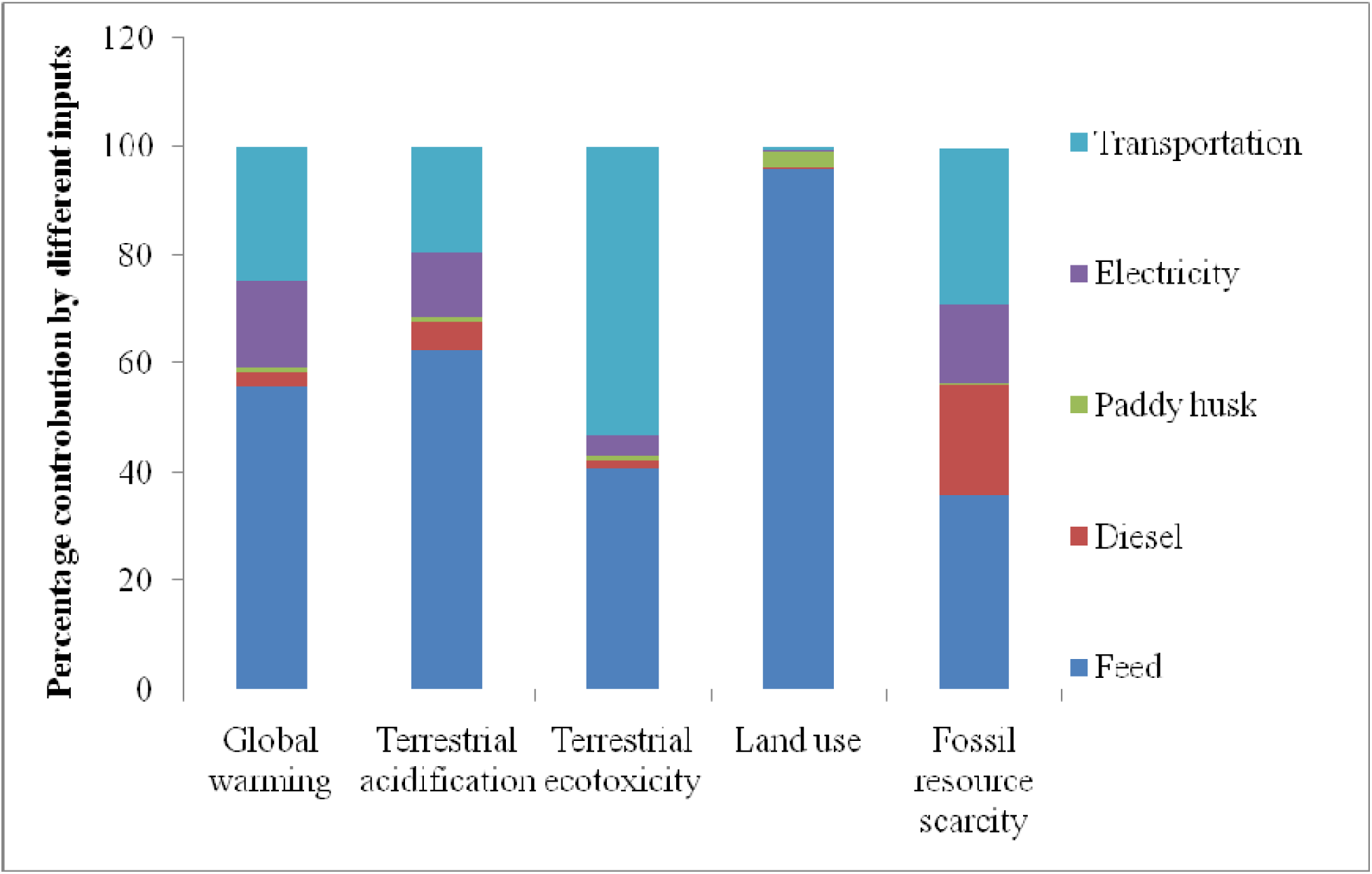
Contribution (%) of various inputs to different impact categories

**Table 5:** The environmental impact of one kg live weight broiler chicken produced.

| Impact category | Reference Unit | Result |
| --- | --- | --- |
| Global warming potential | kg CO<sub>2</sub>-eq | 3.77 |
| Stratospheric ozone depletion | kg CFC11-eq | 0.00002 |
| Ionizing radiation | kBqCo-60-eq | 0.022 |
| Ozone formation, human health | Kg NO<sub>x</sub>-eq | 0.013 |
| Fine particulate matter formation | kg PM2.5-eq | 0.0071 |
| Ozone formation, terrestrial ecosystem | kg NO<sub>x</sub>-eq | 0.013 |
| Terrestrial acidification | kg SO<sub>2</sub>-eq | 0.016 |
| Freshwater eutrophication | kg P-eq-eq. | 0.00028 |
| Marine eutrophication | kg N-eq | 0.0067 |
| Terrestrial ecotoxicity | kg 1,4-DCB | 9.11 |
| Freshwater ecotoxicity | kg 1,4-DCB | 0.012 |
| Marine ecotoxicity | kg 1,4-DCB | 0.013 |
| Human carcinogenic toxicity | kg 1,4-DCB | 0.026 |
| Land use | m<sup>2</sup>a crop-eq | 4.63 |
| Mineral resource scarcity | kg Cu-eq | 0.012 |
| Fossil resource scarcity | kg oil-eq | 1.09 |
| Water consumption | m<sup>3</sup> | 0.047 |

Broiler feed accounted for the greatest share of the environmental impacts of the Indian broiler chicken production system. Therefore, a detailed impact analysis of the major feed ingredients was conducted. Among the different feed ingredients (Table 6), maize was responsible for 80% of the total GWP of the feed (1.68 kg CO_2_-eq per kg of live weight), followed by soybean meal (0.27 kg CO_2_-eq per kg of live weight). In addition, maize was found to contribute 87% and 79.48% of the total terrestrial ecotoxicity and fossil resource scarcity, respectively. In the case of land use, soybean meal and maize accounted for 50.87% and 47.53% of total land use by feed. Soybean meal production contributed 50% to the water consumption of feed, while maize production contributed 22.5%.

**Table 6:** Contribution of different feed ingredients to the environmental impact of the broiler chicken production system.

|  | Unit | Feed | Maize | Palm oil | Soybean meal | Fish meal |
| --- | --- | --- | --- | --- | --- | --- |
| Global warming potential | kg CO<sub>2</sub>-eq | 2.10 | 1.68 | 0.08 | 0.27 | 0.05 |
| Terrestrial acidification | kg SO<sub>2</sub>-eq | 0.01 | 0.008 | 0.0002 | 0.001 | 0.0004 |
| Terrestrial ecotoxicity | kg 1,4-DCB | 3.72 | 3.26 | 0.071 | 0.20 | 0.18 |
| Land use | m<sup>2</sup>a crop-eq | 4.46 | 2.13 | 0.057 | 2.27 | 0.0007 |
| Fossil resource scarcity | kg oil-eq | 0.39 | 0.31 | 0.002 | 0.05 | 0.02 |
| Water consumption | m<sup>3</sup> | 0.04 | 0.009 | 0.001 | 0.02 | 0.006 |

## 4. Discussion

In India, the broiler chicken industry is consistently growing at an annual growth rate of 8-10 percent and is expected to expand further. It contributes significantly to the national GDP and provides employment opportunities to millions of people. This industry is mainly growing due to industrialization, an expanding economy, a growing middle class, and the emergence of vertically integrated poultry companies that benefit from economies of scale. However, as the poultry industry continues to grow, the impact on finite resources and impacts from production processes will also increase. Therefore, it is crucial to maintain high efficiency throughout the supply chain. Despite the industry’s significant contribution to the economy, no LCA study has assessed the environmental impacts of the Indian broiler chicken production system. To the best of the authors’ knowledge, this is the first LCA study reporting the environmental impact of the broiler chicken production system in India.

The production and use of broiler chicken feed are major contributors to environmental impacts across all categories. Specifically, broiler feed production can be considered a hotspot for the environmental impacts of broiler chicken production in India. These findings are consistent with numerous previous studies (Bengtsson and Seddon, 2013; Castanheira et al., 2010; Costantini et al., 2021; Duarte da Silva Lima et al., 2019; González-García et al., 2014; Ogino et al., 2021; Pelletier, 2008; Wiedemann et al., 2017). Feed production was examined in detail to identify the main environmental *hotspots* of the broiler chicken production system in India. Among the various components of broiler chicken feed, maize production contributed the most across all categories, except land use and water consumption. Maize is a nutrient-exhaustive crop and requires large amounts of organic and inorganic fertilizers. Besides, it also requires regular application of herbicides, weedicides, and insecticides. As maize accounted for almost 60% of the feed, its contribution to overall environmental impacts was substantial. In line with our findings, Wiedemann et al. (2017) reported that environmental impact was greatest from wheat, followed by sorghum and soybean meal in Australian chicken meat production. Conversely, some previous studies have identified the protein components (mainly soybean) of the feed as the major contributors to environmental footprints (Ogino et al., 2021). This is especially true for European chicken production systems, which heavily rely on imported soybeans from South America, where land-use change is significant (Costantini et al., 2021; Dekker et al., 2013). In the present study, soybeans’ contribution to environmental impact was significant in the land use and water consumption categories. This may be due to lower soybean productivity and irrigation water use. Costantini et al. (2021) reported that protein feeds have the greatest environmental impacts due to land-use change. In the present study, land-use change was not considered due to insufficient information. Nevertheless, the global warming impact of soybeans accounted for only 12.85% of the total feed impact in the current study. Therefore, based on the findings of the present study, it is recommended to use locally produced grains in broiler chicken feed, as the region produces sufficient quantities of maize and soybeans. The production system for maize and soybeans in this region is a low-input, high-output system that uses only rainwater and organic fertilizers, which may have lower environmental impacts. However, most of the produced grains are transported to other regions for livestock feed production.

As reported by Wiedemann et al. (2017), strategies to reduce the environmental impact of feed include replacing grains used in chicken feed and improving the feed conversion ratio (FCR) through genetic improvement. The previous study reported reductions in greenhouse gas emissions and fresh water consumption of 3-4.5 percent and 5 percent, respectively, with an improvement of 0.1 in FCR. McKay et al. (2000) noted a yearly improvement of 0.02 in FCR over the last 30 years, resulting in a reduction of 0.6 kg feed requirements. By improving FCR, the impact on feed production is reduced. Costantini et al. (2021) reported that the feed conversion ratios (FCR) of poultry strongly influence the environmental impact of the entire supply chain. Therefore, improving FCR in broilers is ranked as the top priority to achieve more sustainable production. Dietary alterations, including reducing crude protein intake and optimizing amino acid balance, can lower FCR and reduce manure emissions (Wiedemann et al., 2016). Also, improved FCR could reduce the need for arable land. This could reduce pressure on water and arable land resources and ultimately lower impacts from broiler chicken production farms. In another study, Leinonen and Williams (2015) reported that a low-protein diet coupled with protease improved the environmental performance of broiler chicken production. Ogino et al. (2021) reported that a low-protein diet, along with crystalline amino acids, reduced nitrogen excretion and consequent ammonia emission from poultry manure. Similarly, the use of specialty feed ingredients (SFI), such as supplemented AA and phytase, significantly reduced the global warming potential, eutrophication, and acidification potentials (Kebreab et al., 2016). Such improvisations are crucial for the sustainability of the broiler chicken industry. However, Tallentire et al. (2017) reported that the degree of flexibility to simultaneously reduce several environmental impact categories differed across regions, due to the different feed ingredients available in each region. Besides, Tallentire et al. (2017) and Costantini et al. (2021) reported that the lower the age at slaughter, the lower the environmental impacts of broiler chickens. In the present study, the slaughter age was 42 days; therefore, reducing it to 30-32 days can be explored in future research.

After feed use, transportation, and electricity consumption accounted for the largest proportions of the global warming potential and other impact categories in the broiler chicken production system. In the present study, feed and day-old chicks were transported over a long distance from the source to the broiler farms. The dependence of broiler farms on external suppliers for feed and chicks, due to the absence of local hatcheries and feed manufacturing facilities, results in long-distance transportation of these inputs, which can be a significant source of environmental impact. As the broiler chicken industry is heavily dependent on concentrate feed, strengthening local feed production and supply chains will reduce the environmental impact of feed transport. Similar findings have been reported by Wiedemann et al. (2017), who reported that the grow-out phase contributed the largest proportion to the environmental impact after feed use. To reduce the environmental impact of transportation, it is recommended to strengthen the local feed production (Dekker et al., 2013; Thévenot et al., 2013). Also, a poultry hatchery should be established in the region to produce chicks and supply them to local farms. In addition, suitable climate-specific housing design, quality equipment, proper ventilation, and installation of solar panels for on-farm energy use can lower the environmental impacts of electricity consumption (Thévenot et al., 2013). Wiedemann et al. (2017) and Ogino et al. (2021) suggested energy production from poultry manure/litter as a way to reduce emissions and lower energy demand from production; however, further analysis is warranted in this area. Therefore, it is essential to employ a multi-criteria approach that considers both environmental impacts and economic considerations to improve the sustainability of broiler chicken production.

The LCA of poultry meat and egg production is well documented in North America and Europe (Costantini et al., 2021). While a precise comparison with studies from other countries is difficult, a broader overview can be obtained and may assist in improving the broiler chicken production system in the future. Hence, a comparison of the global warming potential of the present study with already published studies is presented in Table 7. Although on the higher side, the findings of the present study fall within the range of previous studies.

**Table 7:** Comparison of cradle-to-farm gate climate change impact (GWP) of broiler chicken production systems of different countries published in the literature.

| Country | System boundary | GWP (kg CO<sub>2</sub>-eq) | Reference |
| --- | --- | --- | --- |
| Brazil | Cradle to farm gate | 2.70 | Da Silva Lima et al., 2019 |
| Greece | Cradle to farm gate | 3.92 | Giannenas 2017 |
| Italy | Cradle to farm gate | 3.84 | Cesari et al., 2017 |
| France | Cradle to farm gate | 2.2 to 2.7 | Prudencio da Silva et al., 2014 |
| Brazil | Cradle to farm gate | 1.4 to 2.06 | Prudencio da Silva et al., 2014 |
| European Union | Cradle to farm gate | 3.43 | Leip et al., 2010 |
| UK | Cradle to farm gate | 3.087 | Leinonen et al., 2012 |
| France | Cradle to farm gate | 3.12 | Baumgartner et al., 2008 |
| Australia | Cradle to slaughterhouse | 1.8 to 2.2 | Wiedemann et al., 2017 |

Giannenas et al. (2017) reported that land use and land-use change (LULUC: direct and indirect)-related emissions significantly affected the environmental impacts of the broiler production system. In addition, differences in protein content in the diets and in their production areas and productivity also affect the environmental footprint of a poultry production system (Prudêncio da Silva et al., 2014). Similarly, production systems and slaughter age of broiler chicken had significant effects on environmental footprints. Leinonen et al. (2012) reported that the organic chicken production system has higher acidification and eutrophication potentials than the conventional production system. Ogino et al. (2021) reported that slaughter age was significantly correlated with the acidification potential, eutrophication potential, and energy consumption, but not with greenhouse gas emissions. It should be noted that different production systems have different advantages in terms of animal welfare, meat quality, and ecosystem biodiversity. An organic chicken production system promotes and enhances the health of the agroecosystem, including biodiversity, biological cycles, and soil biological activity. With a focus on holistic health management and a biologically active soil, organic chicken farming strives to improve the health and welfare of the birds, meat quality, and environmental sustainability.

### Actionable point

The current study is the first to use the LCA approach to assess the environmental impacts of India’s broiler chicken production system. However, as with any study, there are limitations discussed in the following section that require further research. Nevertheless, the study’s findings suggest that improvements can be made in broiler chicken feed production and the supply chain, as these processes significantly contribute to environmental impact. González-García et al. (2014) reported that there are obvious advantages associated with resource-use-driven impacts, including reduced transportation, reduced integration of energy-rich feeds, and no land transformation. Promoting the use of locally produced animal feed will have lower environmental impacts (Baumgartner et al., 2008). Moreover, establishing a poultry hatchery in the production region will also reduce the environmental impacts of the broiler chicken production system. The other improvement action may include installing a rooftop solar system to harness solar energy, in addition to improvements in housing design and equipment. Therefore, to reduce its environmental burden, several managerial measures have been proposed, such as the use of organic fertilizers, composting, and incineration. This aspect warrants further research, particularly in humid subtropical climates.

### Limitation of the current study

To examine the environmental impact of the broiler chicken production system in India, the current study used primary and secondary data. The authors collected the primary data through personal visits to broiler farms, while the background data were sourced from the Ecoinvent version 3.0 Life Cycle Assessment (LCA) database. The Ecoinvent database primarily comprises global data, and hence, the closest data points were used to substitute the global data for local representation. However, for energy-intensive processes such as transportation and electricity, the Ecoinvent database is representative of India. Also, poultry manure was not included in the system boundary in the present study; previous studies have reported its significant contribution to acidification and eutrophication potential.

## 5. Conclusion

Chicken meat is a crucial component of human nutrition and is expanding steadily worldwide. In India, chicken accounts for 50 percent of total meat production and is the meat that Indian consumers consume the most. The broiler chicken industry in India is vertically integrated and uses modern technologies. The industry is consistently growing at 8-10 percent annually and is expected to continue growing. Although LCA of broiler meat has been extensively studied globally, no research has focused on the broiler chicken production sector in India until now. The present study assessed the environmental impact of broiler chicken meat in India. The primary causes of adverse environmental effects and resource depletion have been found to be feed production procedures, followed by transport and electricity processes. In the feed, maize production is the main environmental hotspot. The environmental impacts per kg live weight are in agreement with those reported in previous studies. The study estimated a GWP of 3.77 kg CO2-eq per kg of live weight, with a cradle-to-farm-gate system boundary. The environmental impact findings published across countries differ widely, largely due to system limitations, rearing conditions, the geographic location where the study was conducted, and data quality. Nonetheless, there is still room for further reductions in environmental impacts in broiler chicken production in India. Accordingly, the present study proposed actionable steps to reduce further the environmental burdens of broiler meat production, including improved feed production, optimized transportation and electricity use, reduced age at slaughter, and genetic improvement. However, additional investigation is required to ensure these possible advancements, particularly in the Indian context. To increase the sustainability of the broiler chicken business in India, the findings of this study are relevant to both academics and policymakers.

## Acknowledgements

The work was supported by the ICAR-Poultry Seed Project, ICAR Research Complex for NEH Region, Nagaland Centre, Medziphema-797106, India (PIMS Code: OXX01915). The authors sincerely acknowledge the cooperation and support of the broiler farms, which generously provided the primary data for the study. Also, the authors acknowledge the support of The Energy and Resources Institute, Delhi, for collaborating in this study.

## Data availability statement

The data that support the findings of this study are available on request from the corresponding author.

